## Supplementary Tables for "Twelve-month specific IgG response to SARS-CoV-2 receptor-binding domain among COVID-19 convalescent plasma donors in Wuhan"

**Supplemental Table 1 Percentage changes of RBD-IgG titers distribution of COVID-19 convalescent plasma donors at early, middle, and late stages following diagnosis.**

| | **Percentage (%)** | | |
|---|---|---|---|
| | **1-2 months** | **6-7 months** | **11-12 months** |
| 1:2560 | 11.28 | 1.14 | 0.90 |
| 1:1280 | 17.95 | 4.55 | 1.65 |
| 1:640 | 26.67 | 9.47 | 6.16 |
| 1:320 | 16.67 | 26.14 | 18.47 |
| 1:160 | 15.64 | 31.06 | 29.73 |
| 1:80 | 6.41 | 17.05 | 24.32 |
| <1:80 | 5.38 | 10.61 | 18.77 |

**Supplemental Table 2 Percentage changes of RBD-IgG titers in plasma donors with low, moderate, or high titers after a long period of time.** Negative, titers less than 1:80. High (high titers), titers of 1:1280 and ≥ 1:2560. Moderate (moderate titers), titers of 1:320-1:640. Low (low titers), titers of 1:80-1:160.

| | | **% at 10-11 months** | | | |
|---|---|---|---|---|---|
| | | **High** | **Moderate** | **Low** | **Negative** |
| Initial | High (n=70) | 8.57 | 54.29 | 35.71 | 1.43 |
| | Moderate (n=107) | 1.87 | 24.30 | 60.75 | 13.08 |
| | Low (n=60) | 0.00 | 11.67 | 36.67 | 51.67 |
